# Spatial pattern formation as a consequence of Protease Competition

**DOI:** 10.64898/2026.02.14.705881

**Authors:** Priya Chakraborty, Subrata Dey, Ranu Kundu, Malay Banerjee, Sayantari Ghosh

## Abstract

Exploring the emergence of spatio-temporal patterns due to nonlinearities in gene expression is a relatively new development. In this work, we explore the effect of resource constraint on gene regulatory motif from both equilibrium and spatio-temporal standpoint, taking into consideration the degradation class of resource, protease. We have demonstrated that protease-tagged degradation can cause an emergent bistability to form in the system in a steady-state scenario. Instead of a graded linear response in protein synthesis, two Saddle-node bifurcations caused by protease competition provide a switch-like response with hysteresis, where two drastically differing protein concentrations can coexist. We next turn our attention to spatio-temporal analysis: we extend our study for a two-dimensional sheet of cells with diffusible protein molecules and report the stationary patterns.To investigate the reasons behind these non-homogeneous stationary patterns, we investigate the traveling wave solution and observe that a stationary pattern is formed by the traveling wave solution. Considering that proteases play a major role in the regulation and expression of genes in a variety of diseased scenarios, the repercussions of this spatial patterning caused by protease competition can be extensive in gene regulatory systems.

## 1 Introduction

A key focus of the recent developments in synthetic biology has been to address host-circuit interaction and context dependencies for further development and better performance of successful synthetic circuits. These context dependencies, poses major challenges to robust synthetic circuit performance, include factors such as host-cell growth state [1], circuit sensitivity to temperature variations [2], and unprecedented coupling via competition for gene expression resources [3–7]. Resource competition, in general, has been found to increase the nonlinearity associated with a gene regulatory dynamics. Emergent multistability, shifting thresholds, unexpected transitions have been investigated in some recent studies [3, 8, 9], as a result of ribosomal pool sharing. It has been shown that spatio-temporal transient patterns emerge in presence of such resource competition [10]. In another recent work [11], it has been observed that post-transcriptional regulators, like microRNAs, when investigated from a perspective of coordinated response, may give rise to spatial heterogeneity and pattern formation. When these resource competitions give rise to spatio-temporal pattern formation complex cellular mechanisms get triggered, which have major impact on cellular decision-making, development, phenotypic heterogeneity, etc. The seminal work of A. Turing has opened a new field of pattern formation in biological systems by the interaction between two morphogens with the help of the reaction-diffusion (RD) model [12]. The mechanism of pattern formation by positional information [13] and hierarchical morphogen gradient, mechanochemical feedback-based pattern formation [14], chemotaxis and pattern formation [15], are some examples of the diverse mechanisms that work individually or collectively in cellular pattern formation. Examples of pattern formation in biological systems are countless, like, patterning during embryonic development [16], morphogenesis [17], organization of neural networks [18, 19] and patterns on body i.e. in wings of butterflies [20], pigment in fish [21] etc. Again, the traveling wave plays an important role in many biological processes such as wound healing [22], development [23], and tumor invasion [24]. Adler’s work [25] confirmed the *E. coli* bacteria’s ability to form traveling bands through chemotaxis in response to external signals. It is observed that wave propagation in bistable systems can lead to stationary pattern formation [26, 27]. While it is difficult to pinpoint the exact reason for pattern formation in each of these cases, researchers are trying to establish an agglomeration of diverse possibilities with experimental evidence for these pattern formations [28, 29]. The existence of a chemical gradient in an embryo plays a key role in pattern formation in *Drosophila melanogaster* [30, 31]. In a hierarchy, a transition from simple gradient to complex patterning can be seen in giraffe [32]. The growth and development of tissues also affect the pattern formation in embryos [33]. However, the study of pattern formation by gene regulatory networks is mostly unexplored, and in recent times, scientists have focused more, theoretically and experimentally, and reported pattern formation [10, 11, 34, 35], by genetic regulatory motifs, including the ones under resource limitation.

While the limitation of transcriptional, post-transcriptional and translational gene expression resources have been linked with pattern formation in some recent studies [10, 11, 34], another very important resource of gene expression pathway, which has not been investigated with this perspective, are proteases. These are a class of enzymes (which includes Trypsin, Chymotrypsin, Pepsin, Rennin, Cathepsin, Neurolysin, Insulinase, etc.) that help in protein breakdown [36–38], which are present in limited quantity inside animals, plants, fungi, and bacterial cells. Cameron et al. synthetically produced this essential cellular degradation module to control the core bacterial processes and antibacterial targets in the wet-lab which further serves as a research tool to study essential gene function and further in an applied system for antibiotic discovery [39]. The limited availability of this essential degradation machinery inside a cell has been established already [40]. It has been reported that non-explicit coupling between two different proteins through protease sharing introduces nonlinearity in an otherwise monostable network [41]. Though a limited number of studies have explored the emergent behaviors in circuits as a consequence of protease competition, no work has been done on studying the system’s behavior in the collective diffusible cellular environment, as per our knowledge.

As there are several examples of protease-induced regulation, it would be very interesting to study how competition for this cellular resource introduces nonlinearity in the regulatory dynamics in a diffusible environment. This mechanism can be understood through the existence of traveling wave solutions and the speed of propagation, which depends on the diffusivities of the two proteins. Our main goal is to explore whether the rate of change of protein concentration from one level to another can be regulated through the diffusivities of two proteins. In this work, we have focused on a positive regulatory motif under protease limitation and investigated how the bistability mechanism induces non-homogeneous stationary patterns. This article is organized as follows. In Sec. 2, we have elaborated on the model formulation. In Sec. 3, the temporal and the steady state dynamics of the motif are explored for a single cell. In Sec. 4, the spatio-temporal behavior of the motif is explored from a mathematical perspective; in Sec. 5, stationary pattern formation by the motifs is reported for a multicellular diffusible setup. Finally, in Sec. 6, we concluded with some future scope of work.

## 2 Modelling Framework

In this section, we describe the basic modeling methodology and reaction-diffusion coupling for the considered genetic motif. Positive regulation/activation is one of the most frequently recurring genetic constructs in living cells, and it performs some key functionalities of cell physiology. Let us consider, two proteins *U* and *V* that compete for a shared and limited pool of protease *P*, as shown in Fig. 1. In this framework, *P* represents the total concentration of protease within the cell. The binding constant of protein *U* and resource (protease) is *res*_*u*_, similarly, the binding constant of the protein *V* and protease *P* is *res*_*v*_. An inducer *I* is present that induces the production of *U*, and *U* further activates the production of *V* at a maximum rate of *a*. The parameter *k* is the Michaelis constant, or the half-saturation constant for the synthesis of protein *V*, i.e., at *U* = *k*, the concentration of protein *V* will be half of the maximal synthesis rate, i.e., *a/*2. The dilution rate of proteins, *α* is arising from the effect of cell growth. A table containing units of the parameters are given in Appendix. A.

**Fig. 1:**
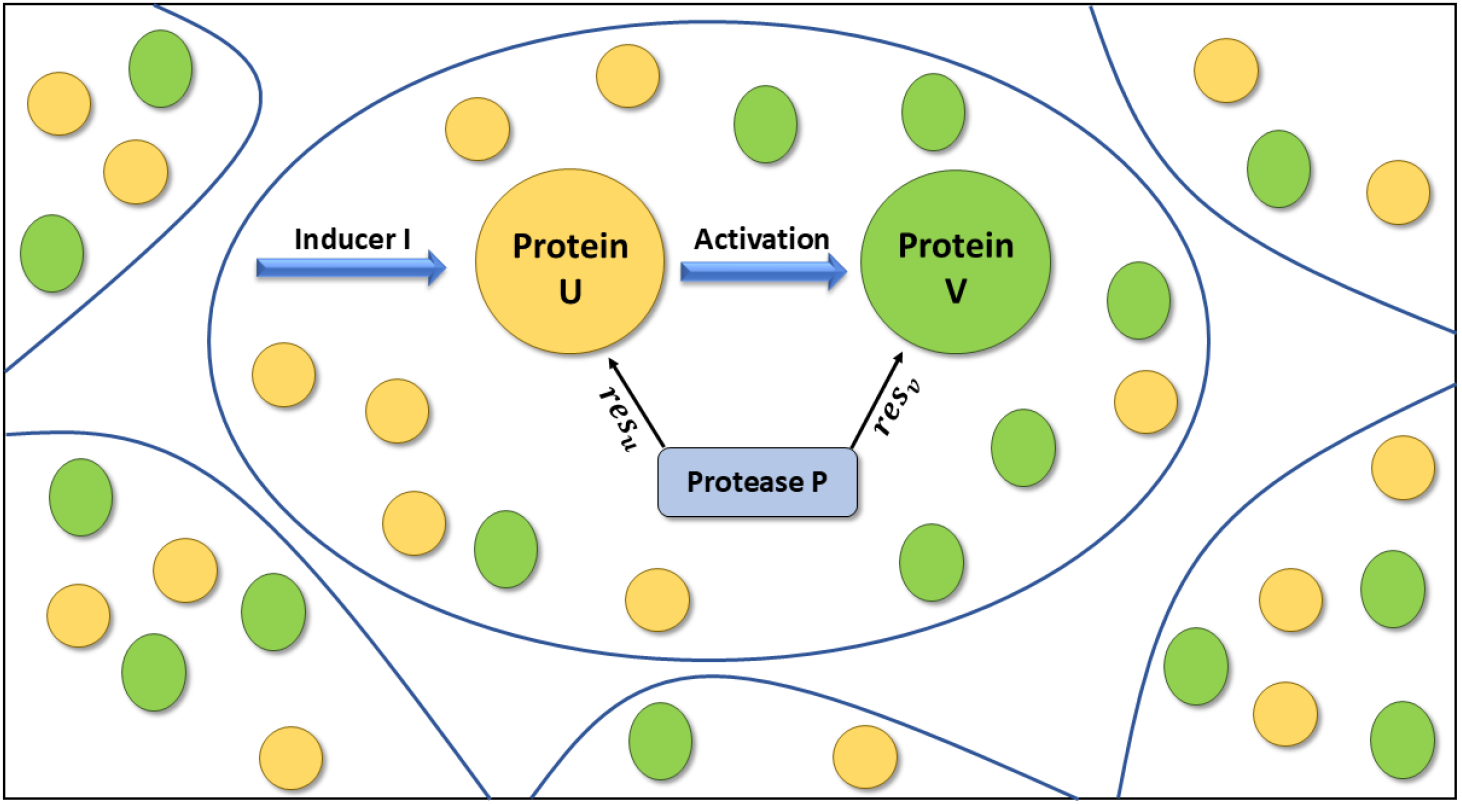
Schematic diagram of the model motif. Cell membranes are shown in blue. A cell is focused in the center, and its intracellular gene regulatory motif of consideration is being demonstrated. Each cell contains the same regulatory motif in the considered two-dimensional array. An inducer *I* induces the protein *U*, and protein *U* further activates the production of protein *V*. Both are tagged with a common pool of protease *P*, which helps in degradation. The binding constant of protease *P* with protein *U* and protein *V* are *res*_*u*_ and *res*_*v*_ respectively. Diffusible molecules of two proteins are represented by corresponding color droplets.

The ordinary differential equations (ODE) representing the dynamics of the above considerations are given by Eq. 1

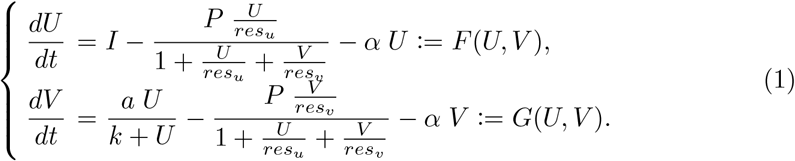

Now, consider a two-dimensional thin sheet of cells in a square lattice Ω (taking into account biological mono-layer tissue formation). Each cell in the lattice contains a single biochemical motif as described before, and we focus on the fundamental process of protein diffusion. Here, in this multicellular arrangement, we consider that the synthesized protein molecules can diffuse across cell membranes. The diffusion coefficient of proteins *U* and *V* be *D*_*u*_ and *D*_*v*_, respectively (here we consider isotropic diffusion, i.e., the diffusion coefficient is the same in both the directions of the considered two-dimensional cell lattice). In Fig. 1, the diffusible protein molecules of two proteins are represented by respective color droplet and the cell membranes are shown in blue boundaries schematically. In a tissue monolayer, the intercellular gaps are negligible and the compact cell arrangement allowed us to study the cell system as a square lattice structure numerically. The set of equations representing the two-dimensional diffusion is given by Eq. 2

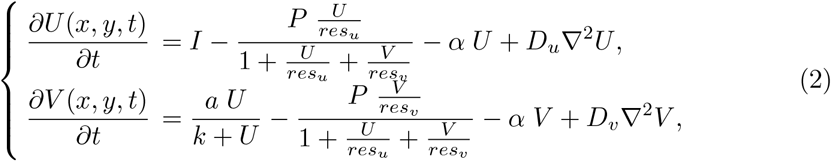

where the ∇^2^ is the Laplacian operator in ℝ^*n*^.

## 3 Steady state behavior of the system

### 3.1 Equilibrium point

Here we discuss the dynamics for the system (1). Let (*U* ^∗^, *V* ^∗^) be the equilibrium point of the system (1) or the homogeneous steady state solution of the system (2), then it satisfy

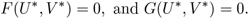

Solving these two equations, we have *U*_∗_ is the roots of the equation

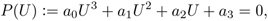

where, *a*_0_ = (*aα res*_*u*_ + *α res*_*v*_*I*) > 0,

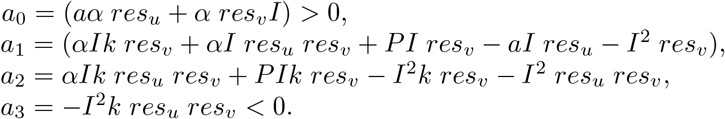

*P* (*U*) is a polynomial of degree 3 with *a*_0_ > 0 and *a*_1_ < 0. Therefore, using Descartes’ rule of signs, *P* (*U*) has at least one equilibrium point and a maximum of three equilibrium points. The system can exhibit saddle-node bifurcation at *I* = *I*_*sn*_, if *P* ^′^(*U*) = *P* (*U*) = 0. We observe that the system is capable of showing two saddle-node bifurcations at 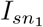 and 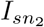, and for 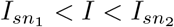, three coexisting equilibria exist. For 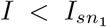 or 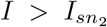, the system has unique coexisting equilibria. In case of three co-existing equilibria, we denote them as 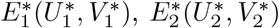 and 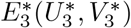 with 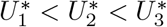.

Considering only the temporal dynamics (i.e., *D*_*u*_ = *D*_*v*_ = 0), the Jacobian matrix of the system at the equilibrium point *E*^∗^(*U* ^∗^, *V* ^∗^) is

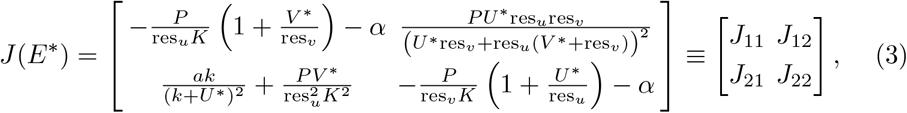

where

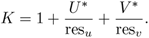

We now provide a rigorous determination of the signs of the diagonal entries of the Jacobian matrix. Recall that

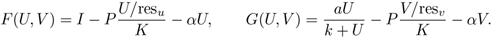

A direct computation yields

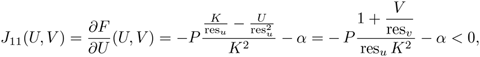

and

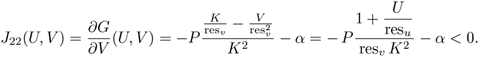

An equilibrium point *E*^∗^(*U* ^∗^, *V* ^∗^) is asymptotically stable (unstable) if both eigen-values have negative (positive) real part and it is a saddle point if the eigenvalues are opposite in sign. In this work, we fix parameter values *res*_*u*_ = 1, *res*_*v*_ = 1, *a* = 90, *k* = 1, *α* = 1 and consider *P* and *I* as bifurcation parameter. In the case of unique equilibrium point, the equilibrium is always asymptotically stable. For the three coexistence equilibrium points, 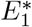 and 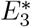 are always asymptotically stable and 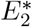 is a saddle point.

### 3.2 Numerical results

#### 3.2.1 Protease sharing introduces bistability in the considered regulatory motif

Protease sharing introduces bistability in the considered otherwise monostable positive regulatory two-gene motif. As we focus on the effects of protease competition, protease regulated degradation of single protein is not a matter of interest here. We equate the rate of changes to zero and explore the steady-state dynamics of the system. In the absence of protease tagged for degradation, the simple positive regulation in protein *V* via protein *U*, which is induced at a constant rate, both the proteins show a graded response with an increase in inducer rate (shown in red curve Fig. 2(a) for protein *U* and Fig. 2(b) for protein *V*). However, sharing the common pool of degradational resource protease makes the dynamics non-linear enough that a binary response can be seen for a range of parameter values for both the proteins (shown in blue curve Fig. 2(a) for protein *U* and Fig. 2(b) for protein *V*). It is a straightforward statement that being tagged by a degradational pool, the concentration of protein will decrease considerably, clear from both Fig. 2(a)-(b) comparing the red and the blue curve; but the emergence of bistability introduces hysteresis in the system via two saddle-node bifurcations. So, the system has a memory of retaining its low/high synthesis state depending upon forward/backward operation. Further, the phase-space plot of protein *U* shows the bistable region (in green), plotted as a function of the total protease available *P* (*x*-axis) and the inducer *I* (*y*-axis), as shown in Fig. 2(c). An extended representation of phase space shows that the bistable region is *closed* in *I* − *P* parameter space (Fig. A.1(b), which demonstrates the possibilities of transition from low response to high response state, without passing through the bistable domain or encountering a discontinuous jump. We have also shown the flow diagram to depict the dynamics with different initial concentrations and to study how the system will reach the steady state in the presence of protease coupling (Fig. 2(d)). The nullcline for proteins *U* and *V* is shown in red and green, respectively. To further assess robustness, we performed a log–fold-change sensitivity analysis of all model parameters, as reported in Fig. A.1(a). This analysis confirms that moderate parameter variations do not eliminate the bistable behavior, although they may shift the boundaries of the bistable region.

**Fig. 2:**
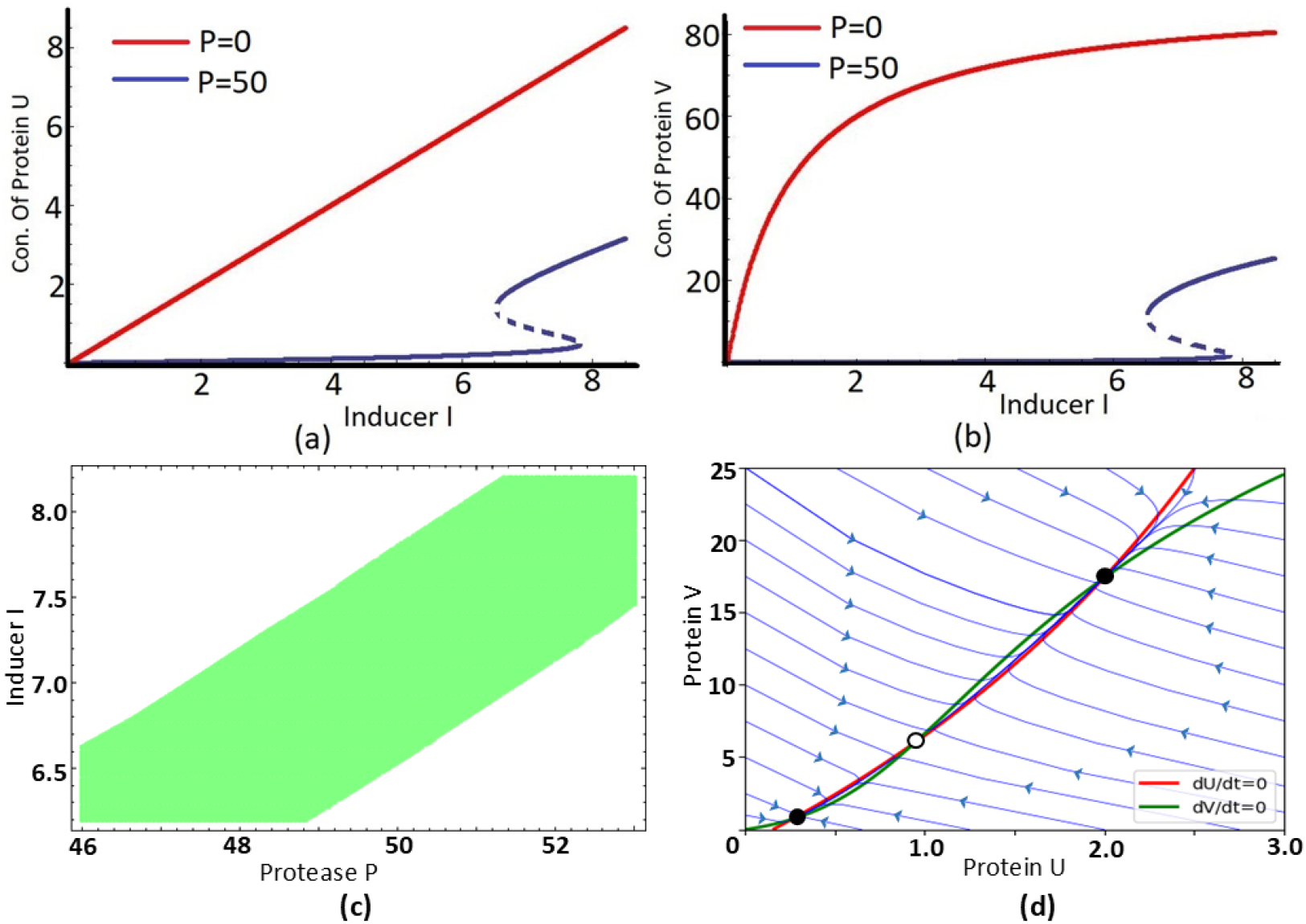
Bifurcation analysis and flow diagram of the motif. (a) Protein *U* shows bistability in presence of coupling with protease, wrt. inducer *I*, shown in the blue curve. Solid lines show stable points and dashed lines show unstable points. In absence of protease coupling the dynamics is monostable, shown in the red solid curve. (b) Protein *V* shows bistability in presence of coupling with protease, wrt. inducer *I*, shown in the blue curve. Solid lines show stable points and dashed lines show unstable points. In absence of protease coupling the dynamics is monostable, shown in the red solid curve. Parameter values for (a) and (b) *res*_*u*_ = 1, *res*_*v*_ = 1, *P* = 50, *a* = 90, *k* = 1, *α* = 1. (c) Phase space of protein *U* for protease *P* vs. inducer *I*. The region of bistability is shown in green, while the monostable region is shown in white. Parameter values here used are *a* = 90 *res*_*u*_ = 1, *res*_*v*_ = 1, *k* = 1, *α* = 1. (d) Phase plane diagram showing trajectories for different initial conditions of protein *U* and *V*. Black solid dots represent stable steady states, and the hollow circle indicates the unstable steady state in the phase space. The blue lines represent the possible trajectories toward a steady state for different initialization in a diffusion-free environment. The red and green curves correspond to the *dU/dt* = 0 & *dV/dt* = 0 nullclines, respectively. Parameter values are *I* = 6.9, *P* = 50, *res*_*u*_ = 1, *res*_*v*_ = 1, *k* = 1 *a* = 90.

#### 3.2.2 Protease affinity regulates bistability for both the proteins

The binding of protein to the protease pool can be asymmetric depending upon several intermediate biological steps. Different biochemical routes of targeting protein substrates by ATP-dependent protease can be further modified by the adapter proteins in the step of binding of substrate to protease [42]. As proteases are enzymes that target and break the peptide bond joining the amino acids together in proteins, it is also biologically relevant to consider that different proteins have different affinities of protease. Here, in this section, we explore the different affinities of degradational resource protease not only affecting its own dynamics but also affecting the dynamics of the other as coupled together. The change in the value of *res*_*u*_ with a fixed value of all other parameters including *res*_*v*_, affects the bistability of protein *U*, along with the change in the bifurcation of protein *V*, as shown in Fig. 3 (a),(c) (Similar thing is shown for a fixed value of *res*_*u*_ including all other parameters with a change in *res*_*v*_, Fig. 3 (b)-(d)). Following our modeling consideration, an increase in (1*/res*_*u*_) effectively increases the degradation of protein *U*. Thus, for a fixed value of inducer *I*, the concentration of *U*, increases with a decrease in 1*/res*_*u*_. The region of bistability, accounting for the robustness of the switch response in the system also changes with the affinity of the resource pool. With a decrease in *res*_*u*_ the bifurcation point of protein *U* shifts to a higher inducer value of *I* along with an increase in the region of bistability (Fig. 3(a)). An increase in region of bistability effectively accounts for a greater switch robustness for the system. As a result of the positive regulation of *U* to *V*, the bistable dynamics of protein *V* reflect the same trend (Fig. 3(c)).

**Fig. 3:**
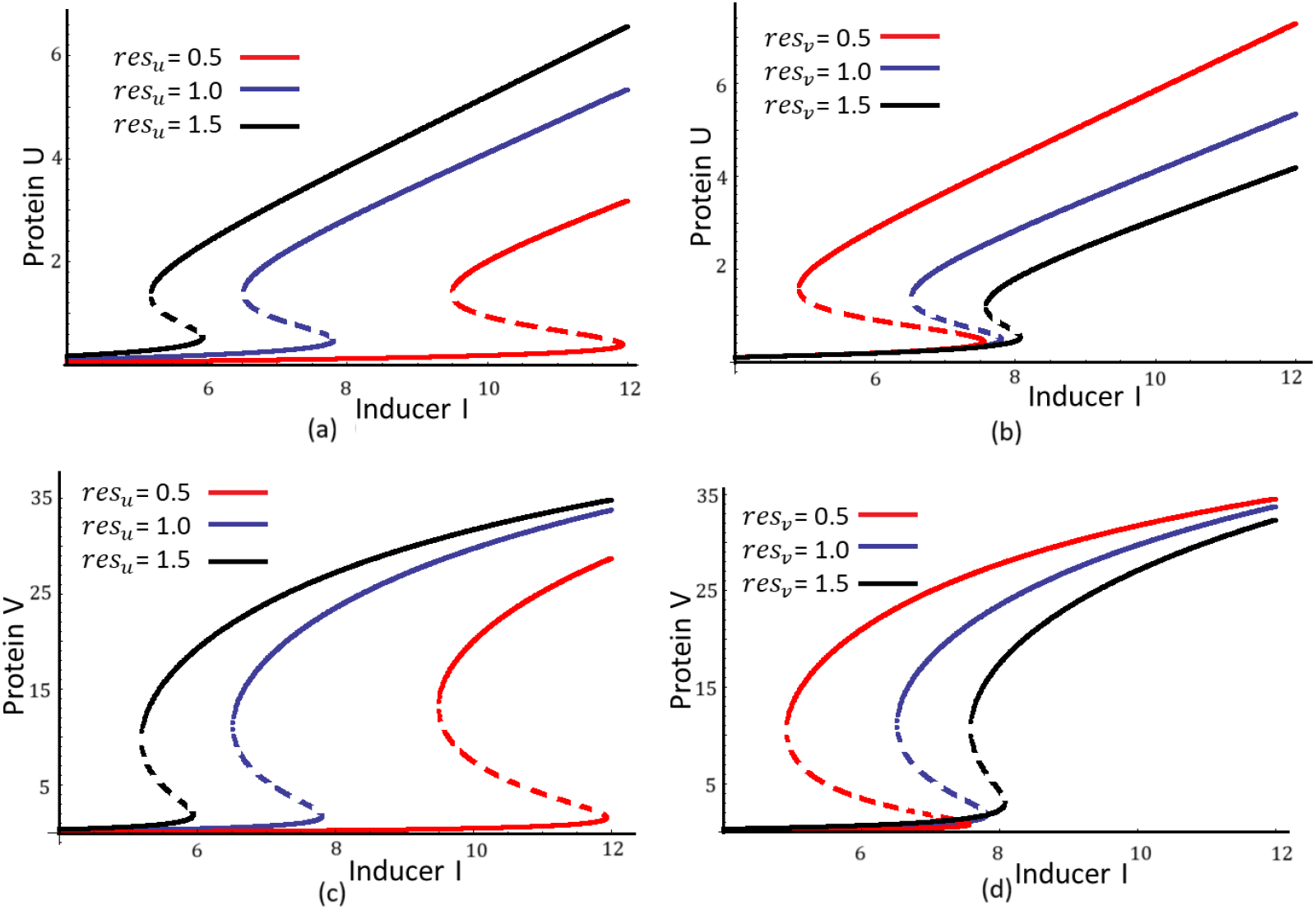
Bistability of both proteins is dependent on protease sharing. (a) Concentration of protein *U* wrt. increase in inducer *I* for three different *res*_*u*_ values 0.5, 1, 1.5 respectively for red, blue, black curve; for a fixed value of *res*_*v*_ = 1. (b) Concentration of protein *U* wrt. increase in inducer *I* for three different *res*_*v*_ values 0.5, 1, 1.5 respectively for red, blue, black curve; for a fixed value of *res*_*u*_ = 1. (c) represent the dynamics of protein *V* at the parameter set of (a), and (d) represent the dynamics of protein *V* at the parameter set of (b). Rest of the parameter values are respectively *P* = 50, *a* = 90, *k* = 1, *α* = 1.

Interestingly, the bistable dynamics of protein *U* is found to be affected by the change in the value of *res*_*v*_, keeping all the other parameters fixed. Here, no direct regulation of protein *U* is being caused by protein *V*. We only consider *res*_*v*_ to capture the effect of the degradation of protein *V* via the protease pool. Thus the change in dynamics of protein *U* via a change in *res*_*v*_ effectively accounts for the coupling of the two proteins via the limited degradational resource pool *P*. From a limited resource pool of *P*, allocating resources more(less) to a particular protein effectively reduces(increases) its availability to the other protein. Thus the dynamics of protein *U*, follow an opposite trend with an increase in *res*_*v*_, keeping all the other parameters fixed in Fig. 3(b), as in Fig. 3(a). Similarly, Fig. 3(c) and Fig. 3(d) can be explained in terms of the protease affinity values of the two proteins.

## 4 Spatio-temporal behaviour of the system

### 4.1 Non-existence of Turing instability

Here, we discuss Turing instability around a coexisting equilibrium point *E*^∗^. Let, 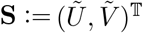 denotes perturbations around the homogeneous equilibrium point *E*^∗^(*U* ^∗^, *V* ^∗^), linearizing about it gives

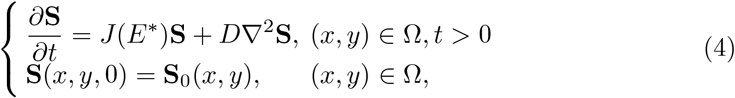

where *D* = *diag* {*D*_*u*_, *D*_*v*_ and *J*(*E*^∗^) is the Jacobian matrix *J* evaluated at the spatially homogeneous equilibrium *E*^∗^.

In Turing instability, we are interested in finding solutions of the linearized system (4) of the form 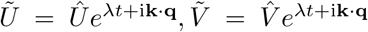, with 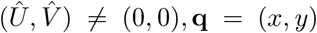, **k** = (*k*_*x*_, *k*_*y*_), and thus get the dispersion relation

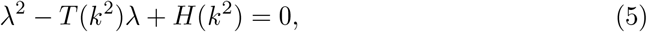

Where 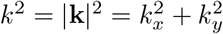, and *T* (*k*^2^) and *H*(*k*^2^) are specified by

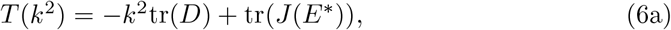

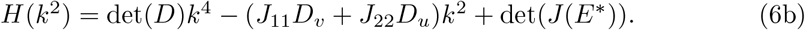

where

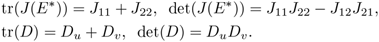

The first assumption for the Turing instability is that *E*^∗^ is locally asymptotically stable for the temporal system, i.e.,

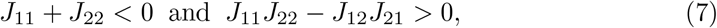

are satisfied.

From tr(*J*(*E*^∗^)) < 0, we obtain *T* (*k*^2^) is negative for all wavenumbers *k*. Therefore, Turing instability can only be obtained in the case of *H*(*k*^2^) < 0 for some non-zero wavenumber *k*. Since, det(*J*(*E*^∗^)) > 0 and det(*D*) > 0, the criteria for *H*(*k*^2^) < 0 becomes

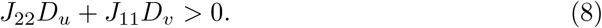

Note that diffusion parameter *D*_*u*_ and *D*_*v*_ are always positive and *J*_11_ + *J*_22_ < 0. Thus, we must have that *J*_11_ and *J*_22_ are of opposite sign. In equation (3), we observe that *J*_11_ and *J*_22_ are always negative for both coexisting equilibrium 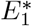 and 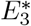. Therefore, the system is unable to produce a Turing pattern solution.

### 4.2 Traveling wave solution

The system doesn’t produce the Turing stationary solution. However, it still shows some stationary solutions under the bistable mechanism for lower diffusion values. To explain the cause of that type of stationary solution, here we discuss the traveling wave solution of the system. First, we consider the system in one-dimensional spatial domain i.e.,

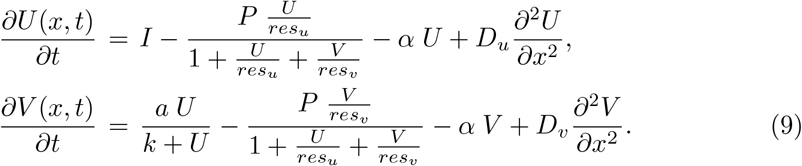

traveling wave solution is a solution of the form

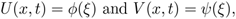

that connects one saddle-node and a stable equilibrium point [43, 44]. Here *ξ* = *x* + *ct* and *c* denotes the wave speed of the traveling wave. For example, in our system, a particular form of traveling wave solution satisfy

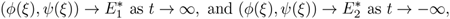

To observe the traveling wave, we consider the following initial conditions in the one-dimensional spatial domain [0,200]

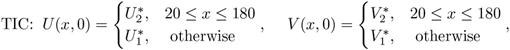

which connects the 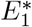 and 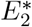. We employed the central difference scheme for the diffusion term and the forward difference scheme for the time derivative in our numerical method. We consider the time step *dt* = 0.01 and spatial step Δ*x* = 1. We fix the parameter values *I* = 6.9, *P* = 50, *res*_*u*_ = 1, *res*_*v*_ = 1, *k* = 1, *a* = 90, *k*_1_ = 1, *k*_2_ = 19, *α* = 1. For *D*_*u*_ = *D*_*v*_ = 10, we observe that the wave propagates as time progresses without changing the wave profile (see Fig. 4(a)). Finally, the solution becomes homogeneous 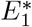 throughout the domain.

**Fig. 4:**
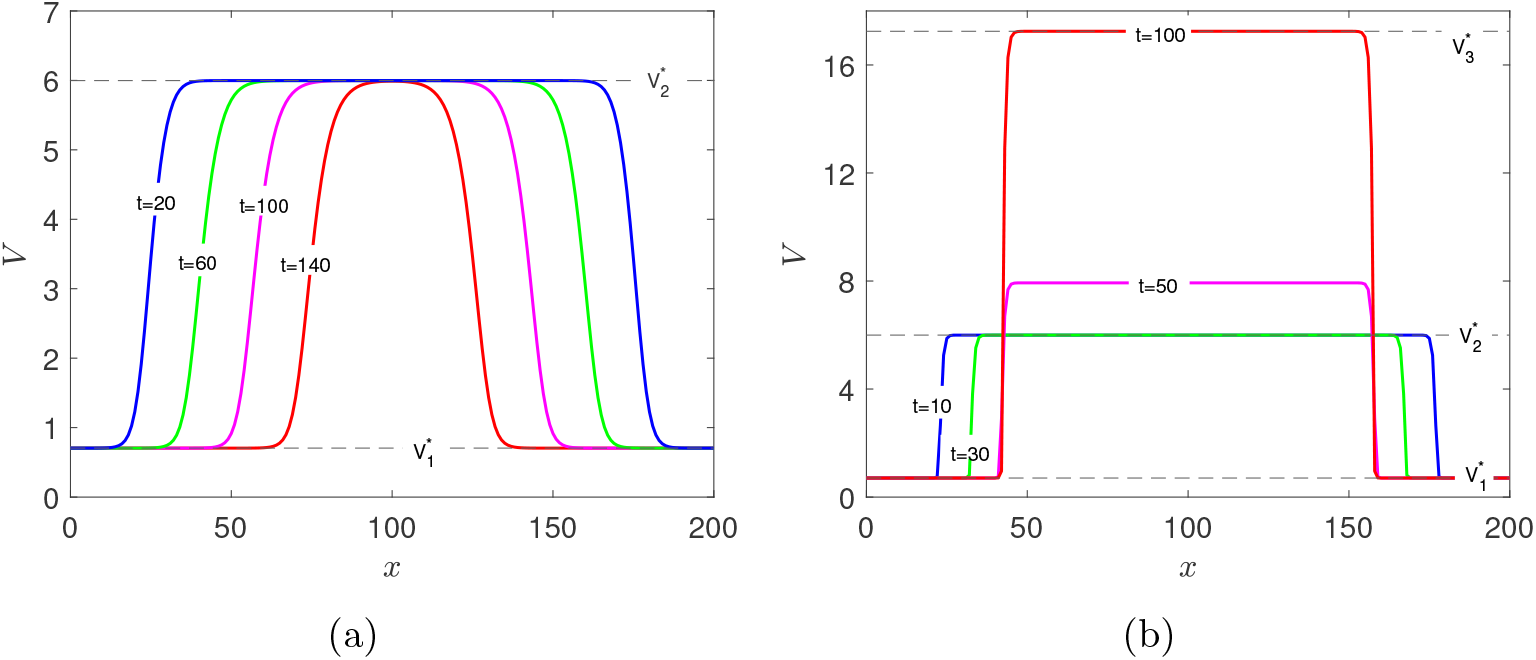
A traveling wave solution with initial conditions connecting 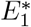 and 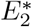 for different values of diffusion coefficients: (a) *D*_*u*_ = *D*_*v*_ = 10, (b) *D*_*u*_ = *D*_*v*_ = 0.1. In (a), the traveling wave propagates smoothly. The colored solutions illustrate the traveling wave at different times: blue (*t* = 20), green (*t* = 60), magenta (*t* = 100), red (*t* = 140). In (b), the traveling wave propagates initially but eventually connects 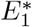 and 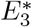. The colored solutions depict the nature of the traveling wave at different times: blue (*t* = 10), green (*t* = 30), magenta (*t* = 50), red (*t* = 100). Here, the black dashed horizontal lines represent the homogeneous steady states, where 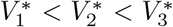. All other parameter values are *I* = 6.9, *P* = 50, *res*_*u*_ = 1, *res*_*v*_ = 1, *k* = 1, *a* = 90, *k*_1_ = 1, *k*_2_ = 19, *α* = 1. and *D*_*u*_ = *D*_*v*_ = 0.1.

Now, if we consider a small value of diffusion, specifically *D*_*u*_ = *D*_*v*_ = 0.1, we observe that the wave does not propagate after some time (in particular after *t* = 33) and instead leads to a stationary solution that connects the two coexisting homogeneous steady states, 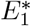 and 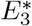 (see Fig. 4(b)). Note that 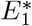 and 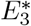 are both stable homogeneous steady states. At a low diffusion rate, the bistability between 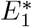 and 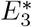 gives rise to a non-homogeneous stationary solution originating from the traveling wave-type initial conditions (TIC).

## 5 Pattern formation

Let us consider a two-dimensional array of (200 × 200) cells where the position of each cell is discretized as *x*_*i*_ where *i* ∈ (1, 200) in *x* directions and *y*_*i*_ where *i* ∈ (1, 200) in *y* direction.

### 5.1 Bistablity induced stationary pattern

We have observed that bistability between 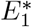 and 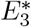 leads to a non-homogeneous stationary solution under the initial condition TIC in one-dimensional spatial domain. Now, we consider similar initial conditions but in two-dimensional spatial domain as follows

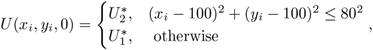

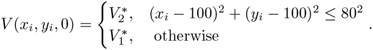

The corresponding surface plot of *V* is shown in Fig. 5. Here, we find that at *t* = 25, *t* = 50, and *t* = 100, the radii of the cylinder-like structure are 120.5, 79, and 78, respectively, while the heights of the cylinder-like structure are 5.99, 8.05, and 17.28, respectively. Therefore, the size of the cylinder-like structure decreases in radius and increases in height, eventually leading to a stationary solution connecting 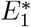 and 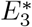. The final solution looks like a rotation of the stationary solution obtained in Fig. 4(b) around *V* axis at the cell (*x*_*i*_, *y*_*i*_) = (100, 100). Under the random initial condition, the same stationary scenario is observed but in an irregular position, which we will discuss next. The color bar lengths of all the spatio-temporal p-color solution are the same as that of *V* as of Fig. 5(d). Thus, the bistability mechanism between the two stable coexisting states leads to the formation of the stationary solution.

**Fig. 5:**
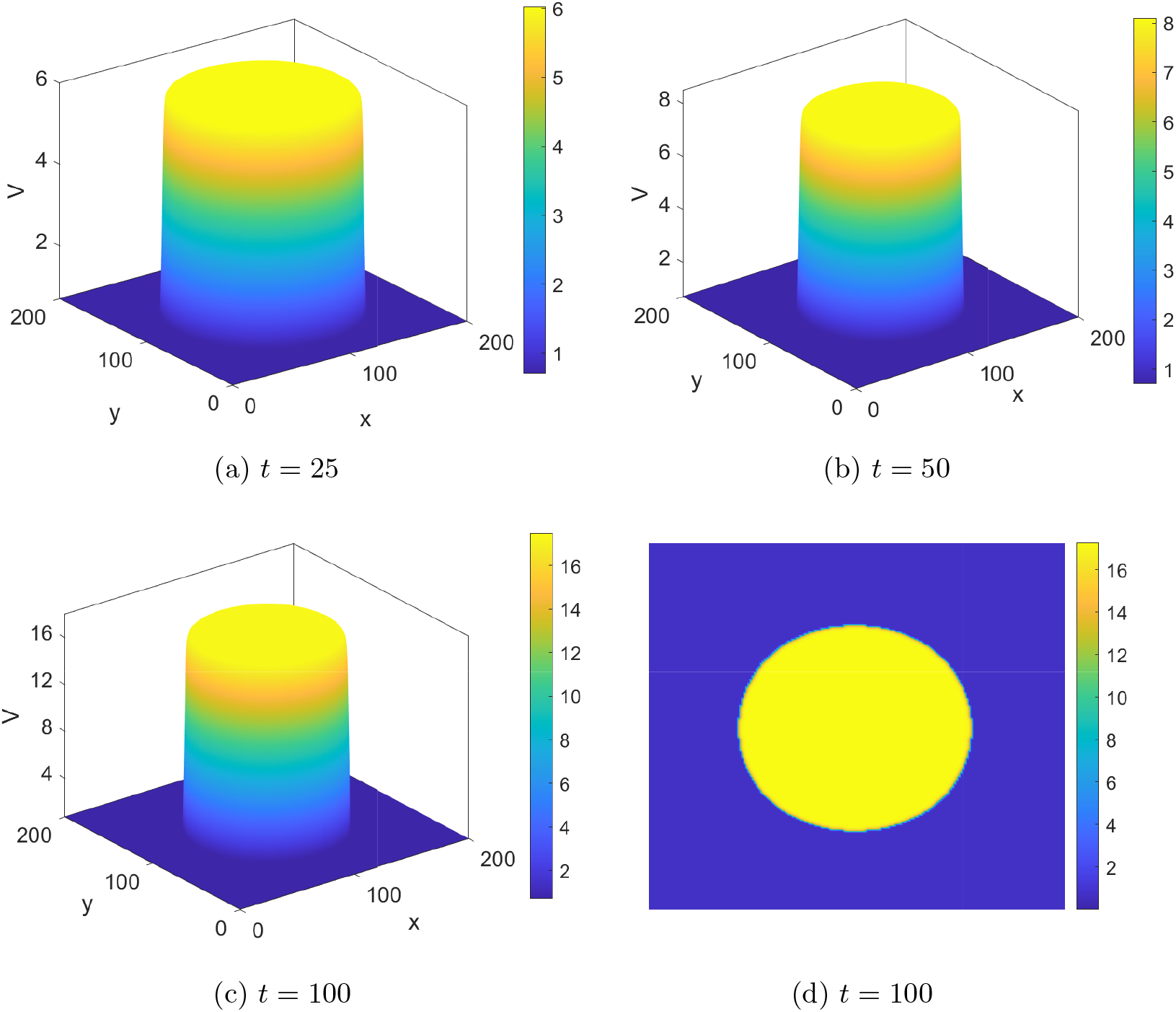
Bistablity induced stationary pattern from invasion traveling wave in two dimension space at different time: (a)-(c) surface plot of *V*, (d) pcolor plot of *V*. Here parameter values are *I* = 6.9, *P* = 50, *res*_*u*_ = 1, *res*_*v*_ = 1, *k* = 1, *a* = 90, *k*_1_ = 1, *k*_2_ = 19, *α* = 1 and *D*_*u*_ = *D*_*v*_ = 0.1.

### 5.2 Stochastic initialisation

We have considered a random initialization following the conditions

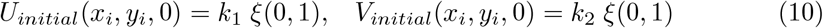

A scaling factor *k*_1_ is randomized via a term *ξ* by picking up a number randomly between 0 to 1 for *U*, and similarly *k*_2_ for *V*. We have tested for different diffusion coefficients as reported in Fig. 6 and reported respective spatio-temporal patterns for protein *V*. To understand the effect of diffusion, in the considered two-dimensional sheet of the cellular array, we increased the diffusion coefficient systematically. For diffusion coefficient of *D*_*u*_ = *D*_*v*_ = 0.01, starting from initial randomization, the system eventually reaches its nearby steady state, but the diffusion is not prominent as no significant change is seen afterward with the increase in time Fig. 6 (a)-(c). For diffusion coefficient 0.05 (Fig. 6(d)-(f)), 0.1 (Fig. 6(g)-(i)), 0.15 (Fig. 6(j)-(l)) respective time evolution and stationary patterns are shown in respective figures. The cells with low protein concentration are shown in color blue, and the cells with high protein concentrations are shown in color red. Starting from randomised initial conditions, first, a binary cell cluster with every cell in one of the two steady states is found. With time, the formation of islands of high and low synthesis states is observed. For different diffusion coefficients, the figures of the 1^*st*^ column are taken after time 10, the figures of 2^*nd*^ column are taken after time 3000, and the figures from the 3^*rd*^ column are taken after time 6000. No effective change in patterns comparing the spatio-temporal distribution of the cellular sheet after time 3000 and 6000 is observed (in each case for a constant diffusion coefficient in individual studies, up to diffusion coefficient *D*_*u*_ = *D*_*v*_ = 0.15). However, it is clear that the island sizes are getting bigger with an increase in diffusion coefficient in the system. Now, for diffusion coefficient 0.2, we can see the pattern is no longer stationary; a transient evolution of spatio-temporal pattern eventually emerges to a fixed steady state (high synthesis state here).

**Fig. 6:**
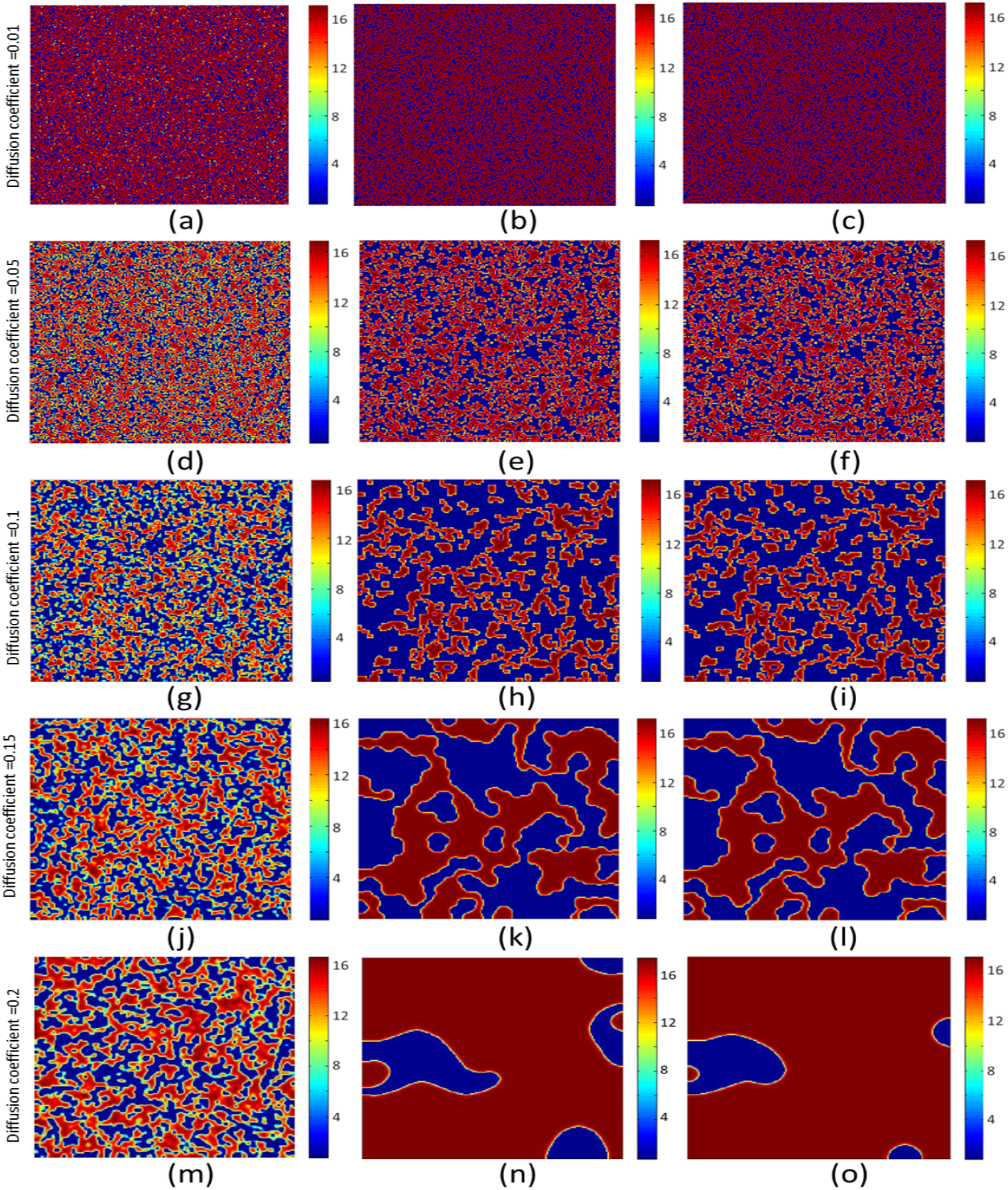
Spatio-temporal pattern for protein *V*, for stochastic initialization. Parameter values are *k*_1_ = 1, *k*_2_ = 19, *I* = 6.9, *P* = 50, *res*_*u*_ = 1, *res*_*v*_ = 1, *k* = 1 *a* = 90. Diffusion coefficients for respective panels are mentioned accordingly. Snapshots are taken accordingly (a),(d),(g),(j),(m) after time 10; (b),(e),(h),(k),(n) after time 3000; (c),(f),(i),(l),(o) after time 6000.

Here, we must note that diffusion increases the instability of the system, while the reaction stabilizes it. Now, we have established that the reaction counterpart of the system is giving a bistable response; due to underlying bistability via the protease-tagged degradation, starting from a stochastic random initial condition, each cell converges to one of its steady states eventually. This can be visualized as convergence through sliding in a hill-and-valley landscape, starting from different initial conditions. Under this circumstances, when diffusion is introduced, it can be considered as random, microscopic movement of the protein molecules. Now in this binary mixture of cell population, the microscopic movement through diffusion eventually help the protein molecules overcome the potential barrier and interconvert from one steady state to another. We know that the depth of the potential well is a function of the reaction rates. Thus depending upon the diffusion coefficient (that effectively supplies instability to the protein molecules and helps overcome the potential barrier) and choice of reaction rates (that sets the depth of potential wells) a balance of reaction-driven stability and diffusion-driven instability makes a stationary pattern in the output. An increase in island size for an increase in the diffusion coefficient shows an increase in conversion from one potential well to the other due to an increase in the microscopic movement in the potential wells. Thus, in (Fig. 6(m)-(o)) the diffusion-driven instability dominates, and the pattern becomes transient in nature.

## 6 Discussion

Spatio-temporal pattern formation caused by nonlinearities of gene expression is a relatively new aspect being explored, in the numerous variety of patterns that can be generated in biological systems. In this paper, we consider a degradation class of resource, protease, the limitation of which is already verified in biological systems. Proteases are known to act as a crucial component in defining the dorsal-ventral pattern of the *Drosophila* embryo [45–48]. The presence of Protease Nexin-1 (PN-1) is found to change the pattern formation during organogenesis and nervous system development in mice [49]. In this paper, we have shown that, in a steady-state scenario, an emergent bistability arises in the system due to protease-tagged degradation for a range of parameter values. It is important to remember that, our model motif does not contain either any feedback loops or cooperativity, the popular driver of bistability in gene expression [50, 51]. However, due to the sharing of a common protease pool between the two proteins, instead of a graded linear response in protein synthesis, bistability in the system is achieved. The green area in Fig. 2(c) represents all (*I, P*) pairs for which two stable steady states coexist, demonstrating that the bistability occurs over a connected and finitely wide region. We found that for no protease sharing, the system is essentially monostable (Fig. 2).

Moving on to spatio-temporal study, we first observe that the system is unable to show Turing pattern formation analytically. Thus, random perturbation around the coexisting equilibrium doesn’t produce any non-homogeneous stationary pattern. But, under the initial conditions (10), the system shows some non-homogeneous stationary patterns. To discuss the mechanism behind this type of pattern formation, we study the traveling solution in bistable system dynamics. Wave propagates as usual for higher diffusive rate of protein (see Fig. 4(a)). However, the traveling wave solution at low diffusive rate of protein leads to a stationary, nonconstant pattern connecting two stable homogeneous steady states. We extend the observation to a two-dimensional sheet of cells. In a large range of scaling for the initial conditions as well as for a large range of monolayer size (tested for 100 × 100, 200 × 200, 500 × 500 monolayer cell arrangement computationally), the pattern is stationary, thus directing towards the robustness of the pattern generated.

There are two striking findings of this work, which we mark as novel contributions: firstly, the patterns we report are not caused by Turing instability, as we have depicted mathematically. Turing instability is a major driver of pattern formation in biological systems, well explored both theoretically and experimentally. However, extreme parametric sensitivity, ad-hoc combination of activator and inhibitor diffusion coefficient and degradation rates, makes it experimentally challenging to reproduce the patterns in synthetic set-ups. Here, we extend our understanding of protease sharing driven bistability in a single cell to a multicellular diffusible setup, and observed this emergent response in form of intriguing patterns. Now, if we focus on the reason behind this pattern formation, it is caused by the underlying bistability of the system. In absence of protease sharing, the cell cluster strictly evolves to a homogeneous spatial state. This driver of pattern formation, in form of protease competition, is another novelty of our work. Conventionally, feedbacks, cooperative binding etc. have been found to drive the nonlinearities in gene regulatory dynamics, causing multistable response. To the best of our knowledge, this is the first study which explores pattern formation driven by protease competition.

To explore the origin of the pattern formation in our system, we reported that the traveling wave propagation can explain the pattern formation in bilayer cell sheets. Very recently, bistability-based pattern formation received the attention of scientists, and synthetic biologists are paying major attention in successfully reconstructing the patterns in lab to avoid difficulties in pattern formation by Turing systems, specially in a system of more than two genes. Under the control of diffusible molecules, Barbier et al. successfully produce spatiotemporal patterns in synthetic toggle switch in *E. Coli* [52]. In a quorum-sensing toggle, the author shows that the circuit exhibits population-wide bistability in a well-mixed liquid environment and generates patterns of differentiation in colonies grown on agar [53].

As we report this emergent bistability-based pattern in the system, we also report that the diffusive rate of protein has a significant role in pattern formation here. Fig. 7 shows that change in diffusive rate causes significant change in the final stationary distribution of two proteins over the sheet of cells. Another important factor that plays its role here is spatial discretization. Biologically, this can be related to the existence of individual cell boundaries. In Fig. A.4 it has been demonstrated the role of this discretization in the observed patterns. This is explained by the fact that moving fronts are a classic example of self-organized wave patterns, representing waves of transition from one stable state to another. A moving front’s velocity and profile are exclusively determined by the medium’s characteristics and are independent of initial conditions. One can typically find either spreading or retreating fronts depending on the properties of a medium. Our results are in agreement with some previous findings [54–58] that report that stationary fronts (or the interface between the regions with two different stable states), as observed in our case, are not observed in continuous media. Only systems with underlying bistability, where a transition from one state to the other occurs, exhibit these stable patterns across a range of parameter values, in the presence of discreteness (such as, non-zero finite cell size). As observed in Fig., the stationary patterns progressively get destroyed and converge into a single state for very large diffusion constants, as the intrinsic length scale of the dynamics becomes much larger compared to the spatial discretization. Thus, if the rate of diffusions are sufficiently weak, or the discretization of space is compatible, the traveling fronts in discrete systems with diffusively connected bistable components may become pinned, resulting in stationary patches. We look forward to the possibilities of novel experimental observations related to our study.

**Fig. 7:**
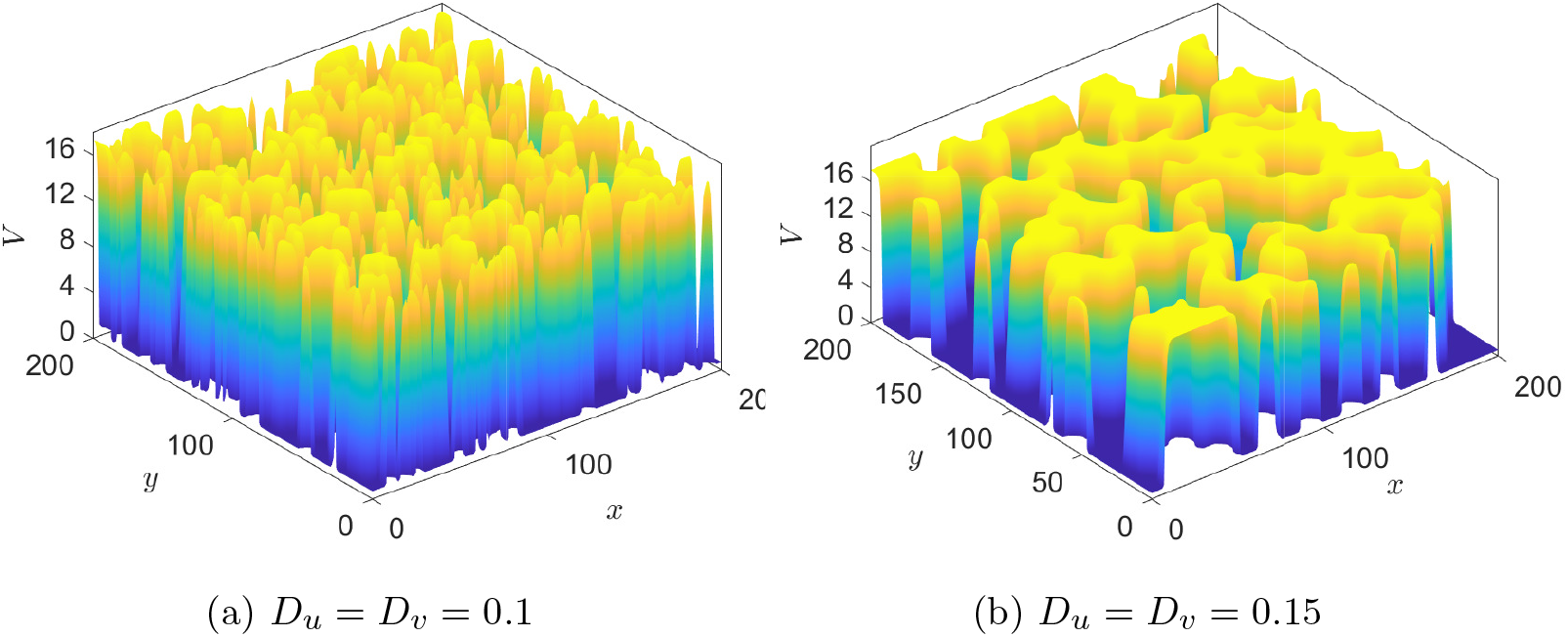
Stationary patterns for different values of diffusive parameter. Here parameter values are *I* = 6.9, *P* = 50, *res*_*u*_ = 1, *res*_*v*_ = 1, *k* = 1, *a* = 90, *k*_1_ = 1, *k*_2_ = 19, and *α* = 1.

Importance of proteases in different regulatory behaviors, including changes in time response in a system [59], stability of oscillatory systems [40], emergent coupling in systems [60, 61] etc., have been noted by several researchers. Thus, the consequences of these spatial patterning observed as a result of protease competition can have far-reaching biological effects. For example, we cannot ignore the role of protease imbalance associated with several diseases. Proteases are an essential class of cellular resources, the imbalance in production, activation, or inhibition of which is majorly related to different diseased scenarios in cells, like cancer, tumors, Alzheimer’s, arthritis, blood clotting disorders, allergies, infections, etc. [62]. Many such disorders have also been related to phenotypic heterogeneity [63–65]. Bistable gene expression has been established as a popular driver of phenotypic heterogeneity [10, 50, 66]. Our results are very interesting from that perspective, as a steady pattern in output can further redirect the cell fate, and these diseased scenarios can be better explored from a perspective of intracellular resource sharing. The dependency of the wavelength of the generated patterns upon initial conditions and diffusion coefficients can also be explored in the future. The system is studied for isotropic diffusion here; however, in real biological systems, diffusion can be direction-dependent, opening further scope of exploration. Moreover, chemical noise, arising from intrinsic stochastic fluctuations in molecular interactions, can influence the stability and behavior of the system. In future study, we plan to extend our system for stochastic reaction-diffusion models and examine for any deviation in system dynamics observed under deterministic scenarios.

## Conflict of Interest

The authors declare that they do not have any known conflicts of interest.

## Acknowledgements

PC and SG acknowledge the support by DST-INSPIRE, India, vide sanction Letter No. DST/INSPIRE/04/2017/002765 dated-13.03.2019.

RK acknowledges the support by DST-INSPIRE, India, vide sanction Letter No. DST/INSPIRE Fellowship/2022/IF220269 dated-22.03.2024.

## Data Availability

The manuscript has no associated data.

**A Appendix**

## System parameters

A detailed explanation of different parameters and their units are given in Table 1. The sensitivity of the system for bistability wrt. system parameters *I, α, P*, *res*_*u*_, *res*_*v*_, *k*, and *a* is reported in Fig. A.1(a). The value of system parameters is varied log fold 0.6 wrt. the reference value as given in the figure caption. The region of bistability is marked green for a range of related log fold change.

**Table 1:**
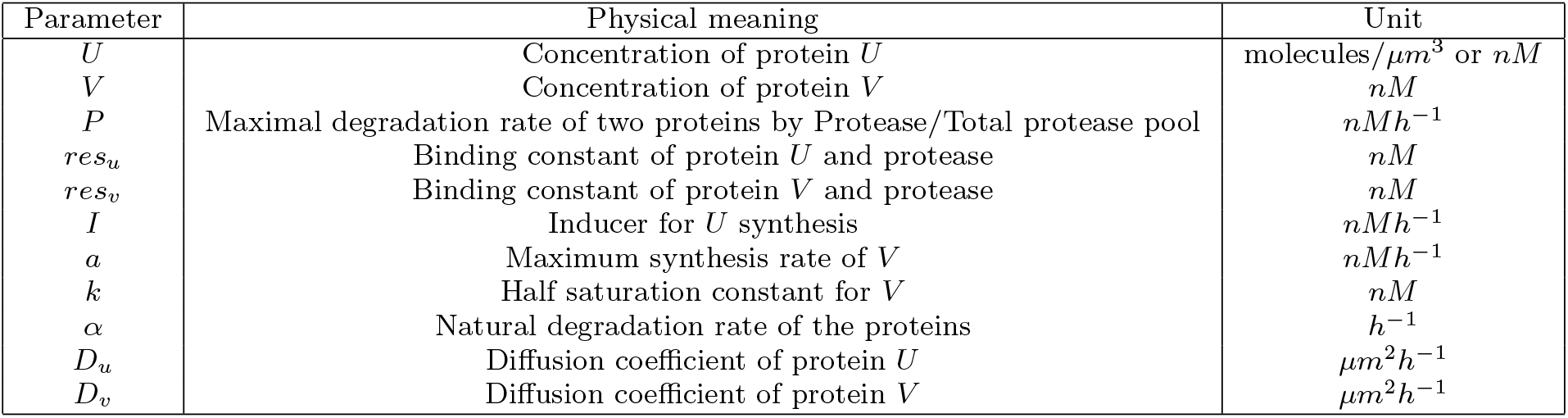
Different parameters, their physical meaning and unit.

**Fig. A.1:**
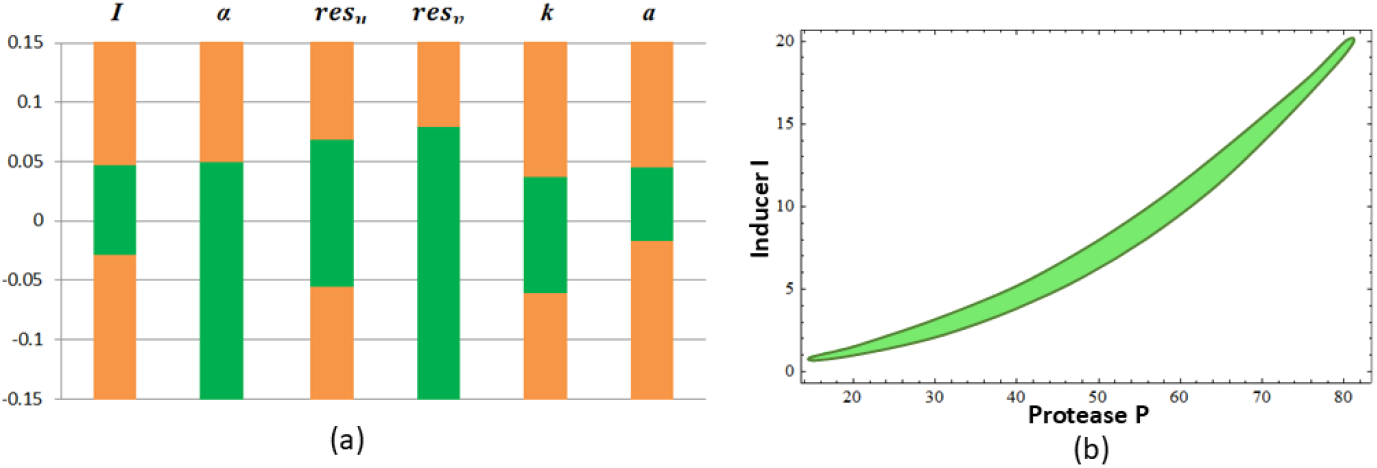
(a) Log-Fold Change sensitivity of different model parameters. Region of bistability is shaded green. Parameter values are *I* = 7, *α* = 1, *res*_*u*_ = 1, *res*_*v*_ = 1, *k* = 1, *a* = 90. (b) Extended representation in phase space shows closed bistable region in *I* − *P* parameter space. Bistable region is shown in color green. Parameter values are *α* = 1, *res*_*u*_ = 1, *res*_*v*_ = 1, *k* = 1, *a* = 90.

## The final stationary pattern & initial randomness

Here, it is interesting to note that the visualization of the final stationary pattern depends on the initial randomization of the system. For two different random initial conditions, *IC*_1_ (Fig. A.2(a) Set I) and *IC*_2_ (Fig. A.2 for Set II), following the criteria as Eq. 9, let us consider that we get two final frames at 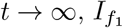 (Fig. A.2(c) Set I) and 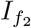 (Fig. A.2(f) Set I) which are both (200 × 200) two-dimensional arrays. Now, if we define pixel wise Absolute Error (AE) as Eq. 11

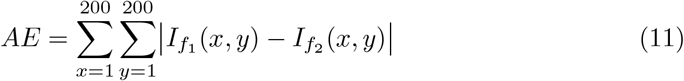

then, we find that the Average Absolute Error (*AE*_*av*_) over 100 consecutive final states *AE*_*av*_ >> 0, which implies substantially different positional information in the final state. However, if we define normalized histograms corresponding to 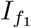 and 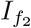 as ℋ_1_(*n*) and ℋ_2_(*n*), the KL divergence,

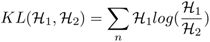

and take its average over 100 consecutive final states, *KL*_*av*_ gives a value very close to zero; i.e.,

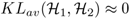

**Fig. A.2:**
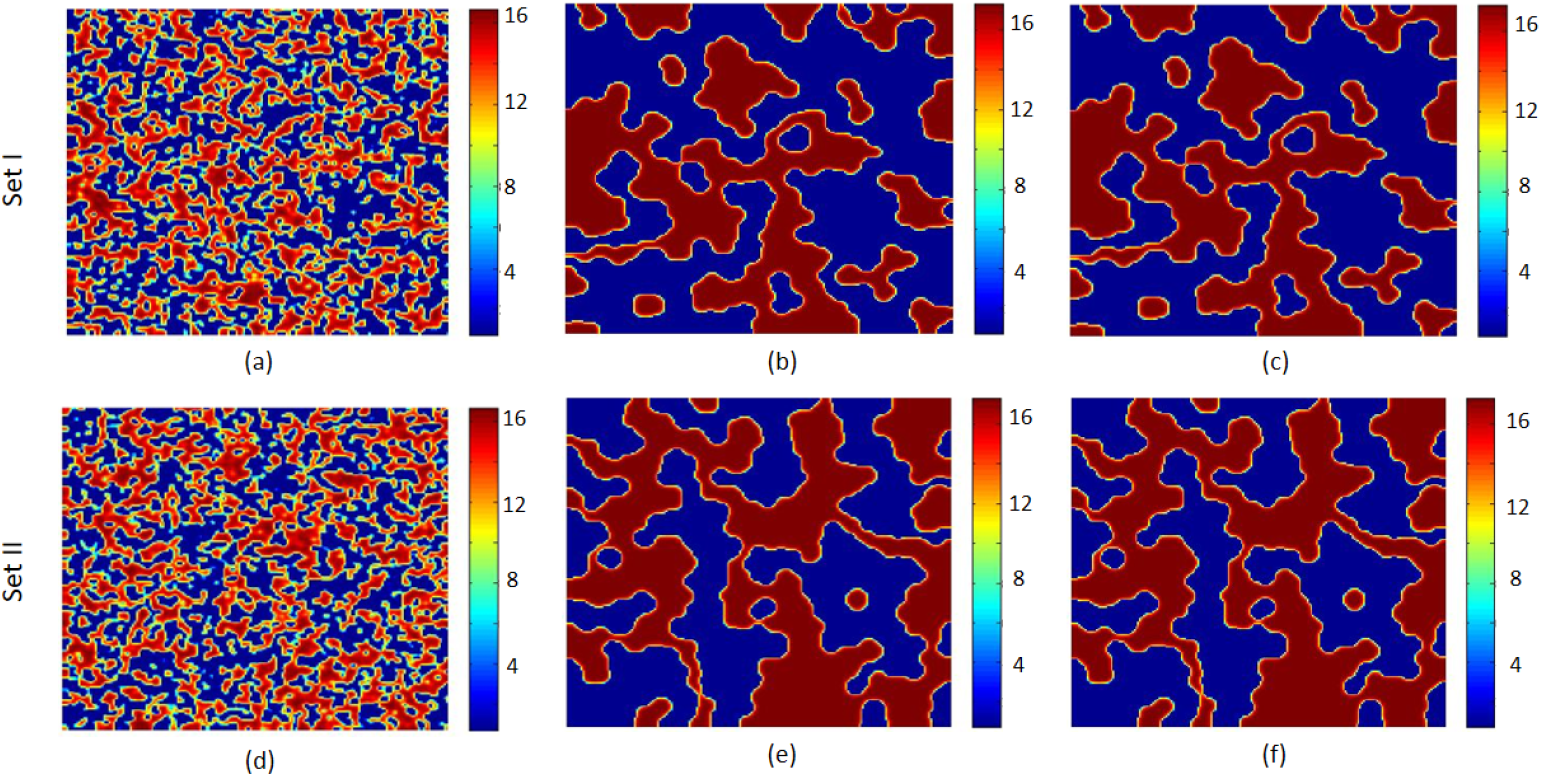
The final stationary pattern is sensitive to the initial randomness of the system. Spatio-temporal pattern for protein *V*, for stochastic initialization of two different sets for the exact same system parameter values. Parameter values are *k*_1_ = 1, *k*_2_ = 19, *I* = 6.9, *P* = 50, *res*_*u*_ = 1, *res*_*v*_ = 1, *k* = 1 *a* = 90, *D*_*u*_ = *D*_*v*_ = 0.15 for both the set of figures.

Thus, we conclude that starting from the same parameter value, step size of iteration, and diffusion coefficients, two different sets of random initialization, the final state of the system accordingly achieves two visually distinct stationary patterns in the output. However, the distribution of total number of cells in the two steady states are almost invariant in the final state.

## Stationary protein patterns as a function of diffusion coefficents

We have explored the effect of the asymmetric diffusion coefficient in the spatiotemporal evolution of the system. For this, we first chose the stochastic random initial state in the lattice by Eq. 10 and then simulated the same randomized lattice sheet for two different sets of diffusion coefficients. Focusing on the spatiotemporal evolution of protein *V*, we see that after equal time, the system with a comparatively higher diffusion coefficient of protein *V* (*D*_*v*_ > *D*_*u*_), shows spatial dominance of higher concentration of protein *V* in (Fig.A.3 Set II), compared to (Fig.A.3 Set I) where (*D*_*u*_ < *D*_*v*_).

**Fig. A.3:**
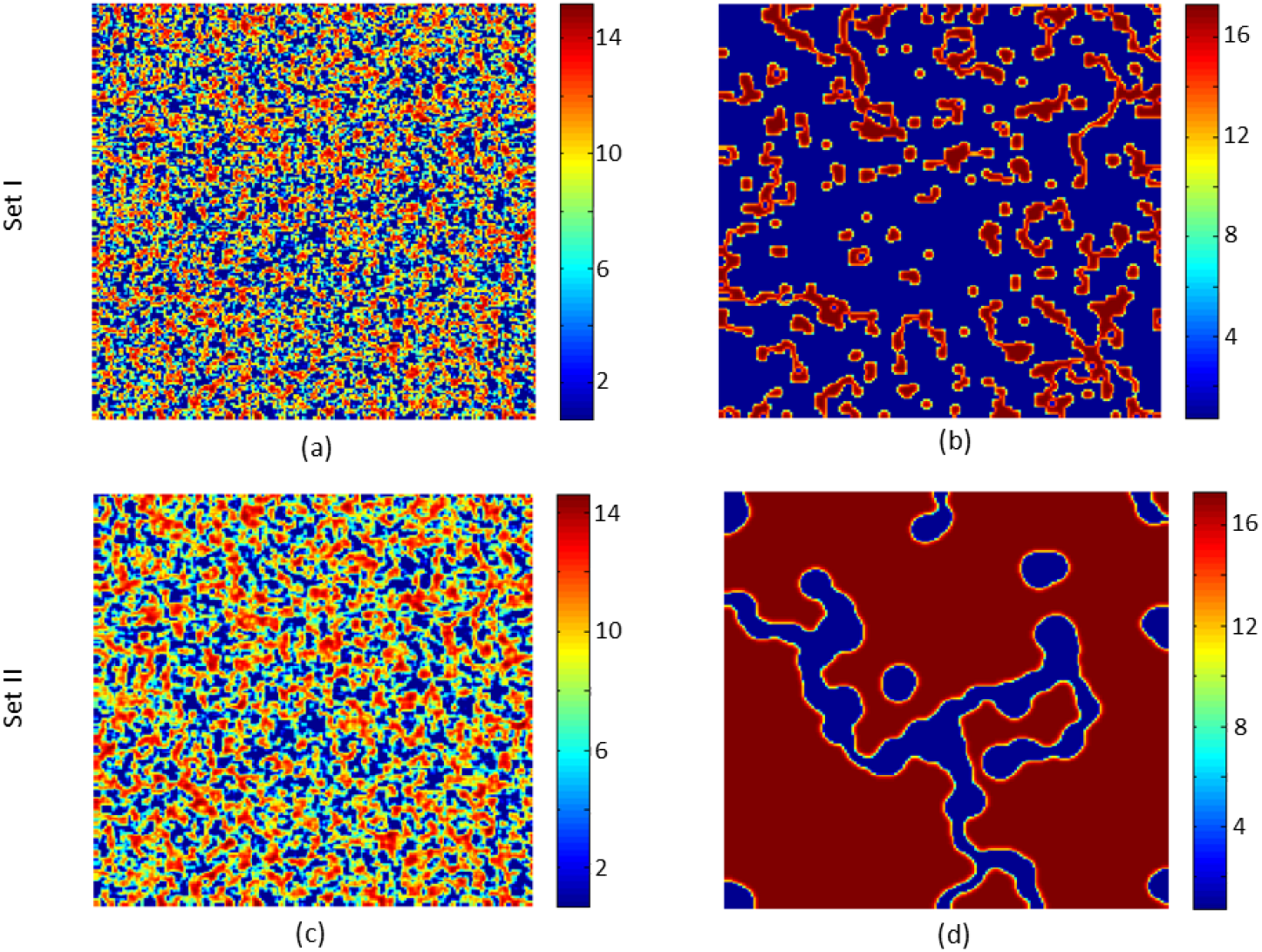
The final stationary pattern is governed by the asymmetric diffusion coefficients of the systems. Spatiotemporal patterns for protein *V* are shown for stochastic initialization under two distinct diffusion coefficient sets: (set I *D*_*u*_ = 0.2, *D*_*v*_ = 0.05 and set II *D*_*u*_ = 0.05, *D*_*v*_ = 0.2). Other parameter values are *k*_1_ = 1, *k*_2_ = 19, *I* = 6.9, *P* = 50, *res*_*u*_ = 1, *res*_*v*_ = 1, *k* = 1, *a* = 90 for both the set of figures. Snapshots are taken accordingly (a),(c) after time 5; (b),(d) after time 3000.

## Effect of spatial discretization on stationary pattern formation

We further observe that the emergence of stationary patterns depends not only on the diffusion coefficients but also on the quantitative discreteness of the spatial scale. Thus, we have also studied the dependency of the stationary patterns on the quantitative discreteness of the spatial scale. As individual cell boundaries (caused by cellular membranes) introduce discretization to this particular system, spatial discretization plays an important role in this dynamics.

In the main analysis [Fig. 6], we reported stationary patterns for diffusion coefficients *D*_*u*_ = *D*_*v*_ = 0.05, 0.1, and 0.15, and transient behavior for *D*_*u*_ = *D*_*v*_ = 0.2, where the spatial separation was set to Δ*x* = 1.0. To assess the impact of spatial resolution, we repeated the simulations with a smaller discretization, Δ*x* = 0.8. To demonstrate this, we repeated the simulations with Δ*x* = 0.8 and reported in Fig. A.4. While stationary pattern is obtained for *D* = *D*_*u*_ = *D*_*v*_ = 0.075 [Fig. A.4(d-f)], gradually for stronger diffusion, extremely slow destabilization start occurring (*D* = *D*_*u*_ = *D*_*v*_ = 0.1) [Fig. A.4(g-i)]. Interestingly, we observed stationary islands for the same *D* = 0.1 for Δ*x* = 1 [Fig. A.4(a-c)]. Finally we observe that the system exhibits transient behavior for *D* = 0.15 [Fig. A.4(j-l)], converging to a homogeneous steady state with time. These findings demonstrate that the persistence of stationary patterns depend jointly on the diffusion strength and the spatial length scale of the system.

**Fig. A.4:**
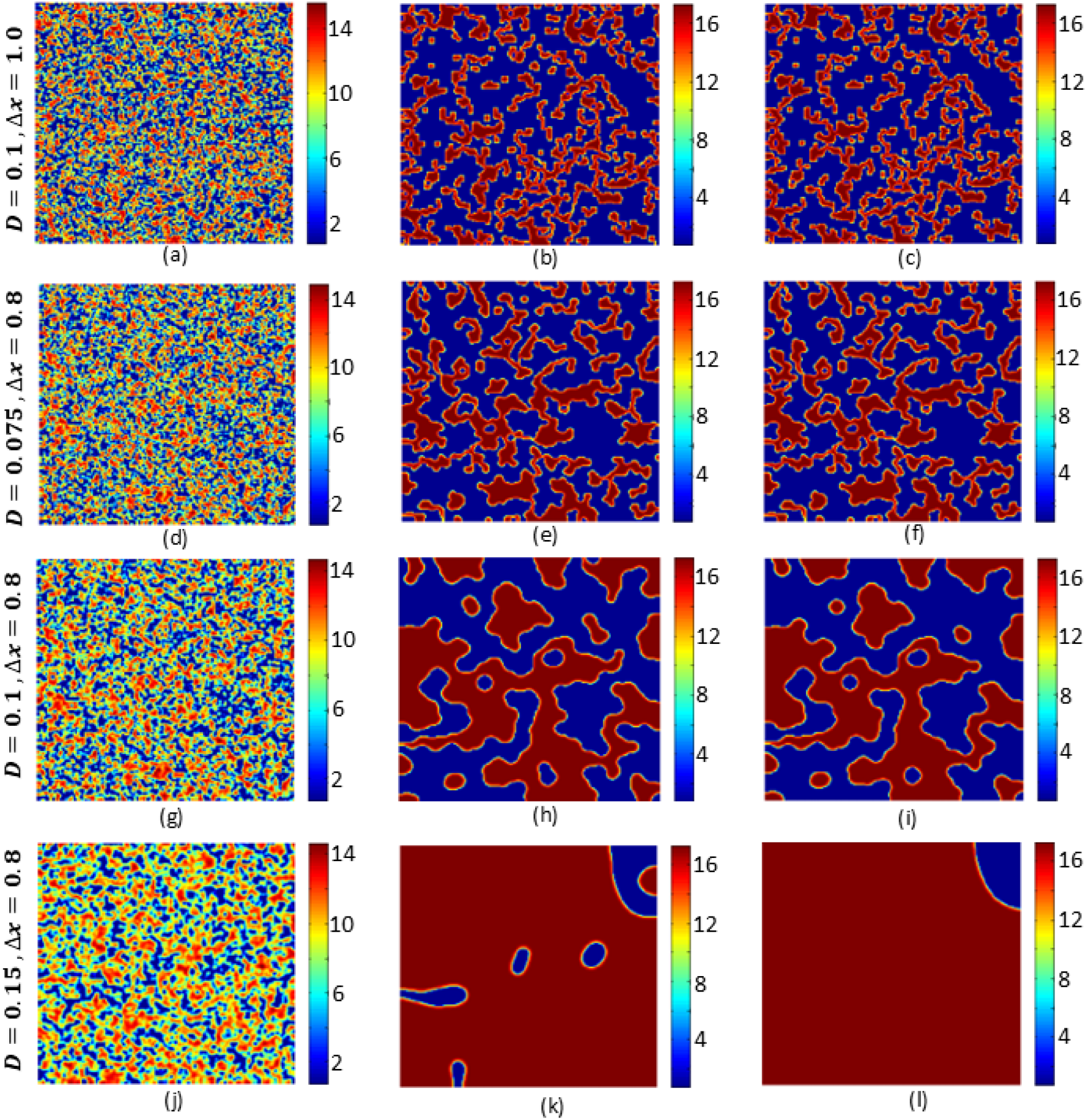
Spatio-temporal pattern for protein *V*, for stochastic initialization at different spatial length scale. Other parameter values are same as Fig.6. Diffusion coefficients and spatial discretization (Δ*x*) for respective panels are mentioned accordingly. Snapshots are taken accordingly (a),(d),(g),(j) after time 5; (b),(e),(h),(k) after time 1000; (c),(f),(i),(l) after time 2000.

## Notes

### Competing Interest Statement

The authors have declared no competing interest.

